# Machine Learning Biogeographic Processes from Biotic Patterns: A New Trait-Dependent Dispersal and Diversification Model with Model-Choice By Simulation-Trained Discriminant Analysis

**DOI:** 10.1101/021303

**Authors:** Jeet Sukumaran, Evan P. Economo, L. Lacey Knowles

## Abstract

Current statistical biogeographical analysis methods are limited in the ways ecology can be related to the processes of diversification and geographical range evolution, requiring conflation of geography and ecology, and/or assuming ecologies that are uniform across all lineages and invariant in time. This precludes the possibility of studying a broad class of macro-evolutionary biogeographical theories that relate geographical and species histories through iineage-specific ecological and evolutionary dynamics, such as taxon cycle theory. Here we present a new model that generates phylogenies under a complex of superpositioned geographical range evolution, trait evolution, and diversification processes that can communicate with each other. This means that, under our model, the diversification and transition of the states of a lineage through geographical space is separate from, yet is conditional on, the state of the lineage in trait space (or vice versa). We present a likelihood-free method of inference under our model using discriminant analysis of principal components of summary statistics calculated on phylogenies, with the discriminant functions trained on data generated by simulations under our model. This approach of model classification is shown to be efficient, robust, and performant over a broad range of parameter space defined by the relative rates of dispersal, trait evolution, and diversification processes. We apply our method to a case study of the taxon cycle, i.e. testing for habitat and trophic-level constraints in the dispersal regimes of the Wallacean avifaunal radiation.

## INTRODUCTION

Model-based historical biogeography methods have advanced remarkably in recent years, facilitating the statistical investigations of a broad range of questions that relate geographical history to the evolutionary history of lineages. The “Dispersal-Extinction-Cladogenesis” (DEC) model was a crucial contribution to the field, providing a rigorous probabilistic framework to model dispersal and vicariance processes informed by phylogenetic and geographical data (Ree et al. 2005; Ree and Smith 2008). Under this DEC model, numerous studies into the geographical origins of groups as well as the histories of changes to the geographical ranges of lineages in those groups over time have been carried out (e.g., de Bruyn et al. 2014; Loader et al. 2014; Mitchell et al. 2014). Powerful and flexible as the DEC model was in answering questions relating to the geographical history of lineages, it was limited in trying to answer questions relating to the process of dispersal itself. The original DEC model treated dispersal as a process that was uniform across lineages. Matzke (2014) extended the original DEC model to incorporate founder-event “jump” dispersal, and its “`BioGeoBears`” implementation allowed for statistical selection of the types of dispersal processes operating in a system using information-theoretic approaches. However, even with this, the fundamental assumption of the DEC framework that all lineages are exchangeable with respect to the dispersal (as well as other) processes remained. With this constraint, studies trying relating ecological processes to phylogenetic, macro-evolutionary, and biogeographical process, or trying to understand how differences in lineage biologies and ecologies contribute to differences in biogeographic patterns, were precluded.

A major advance in integrating ecology with historical biogeography was the geographic state speciation and extinction model (“GeoSSE”; Goldberg et al. 2011), which related geographic distribution to rates of diversification in a statistical probabilistic framework. By using geographic areas occupied by a species as proxies for habitat preferences, the GeoSSE approach provided for the ability to study the relationships between ecology and diversity. Naturally, this approach was restricted by only being able to consider cases in which geographical area could, in fact, be taken as a proxy for habitat (as opposed to being able to model ecology and geography as separate, even if reciprocally interacting, classes of phenomena). But, in addition, it still treated all lineages as exchangeable with respect to the dispersal process, and thus did not allow the modeling of ecology (whether by geographical proxy or not) on dispersal, and thus the effect of lineage ecology on geographical range evolution. Furthermore, the ecologies of the lineages themselves were considered to be fixed in evolutionary time, instead of evolving along with their geographies.

This field of inquiry, i.e., the study of the relationship between ecology and their effects on patterns of geographic distribution of lineages, is of long-standing interest in biogeography. For example, the taxon cycle hypothesis (Wilson 1961; Ricklefs and Bermingham 2002) proposes linkages between trait-evolution, niche shifts, dispersal, and diversification dynamics. Taxon cycle theory posts a particular combination of ecological and evolutionary constraints that lead to a cyclical pattern of biogeographic expansion and contraction associated with niche shifts and diversification. Habitat-dependent dispersal is a necessary (but not sufficient) component of the taxon cycle as well as other biogeographic hypotheses. Such habitat-dependency may be implied by correlations between range size/endemism and habitat type. For example, many studies have found statistical support for correlations between habitat affinity and the species range extent, implying habitat-dependent dispersal (e.g., Ricklefs and Cox 1972, 1978; Dexter 2010; Economo and Sarnat 2012; Carstensen et al. 2012). However, such correlations are potentially misleading about habitat-driven dispersal in the context of the other processes shaping biogeographic patterns such speciation, extinction, and niche evolution.

Our goal is to develop a statistical, process-based model-choice framework to determine if we can discriminate habitat-constrained dispersal regimes from cases where dispersal is not constrained by habitat. More generally, we investigate whether we can identify cases where a particular ecological trait does indeed regulate dispersal of lineages throughout the history of a group. To address this goal, we need an approach that allows for separate geographical as well as ecological state spaces, and conditions transitions of lineages through the geographical state space (which includes range gain through dispersal as well as range loss through extirpation) on the state of the given lineage in ecological space. In this paper we first introduce a continuous-time discrete-area model that not only incorporates diversification processes (i.e., lineage birth and death) and biogeographical processes (e.g., dispersal) as submodels, but also a trait evolution submodel that informs the diversification and biogeographical processes. We then describe a likelihood-free approach for carrying out inference under this model, using simulation-trained multivariate discriminant analysis classification. As a test case, we apply our approach to the Wallacean island bird radiations; a systems in which taxon cycles have been proposed and, more specifically, in which habitat-constrained dispersal has been suggested as an important mechanism in regulating their radiation (Carstensen et al. 2012).

## MATERIALS AND METHODS

### “Archipelago”: A Trait-Driven Biogeographical Phylogenesis Model

The “Archipelago” model generates spatially-explicit phylogenies under a complex of superpositioned stochastic processes, in which the per-lineage speciation, extinction, and dispersal events each occur under independent stochastic processes. The rates of speciation, extinction, and dispersal are not (neccessarily) homogenous, but instead are determined by the states of traits that are associated with the given lineage. These trait states (which can represent any aspect of a lineage: biological, ecological, etc.) are, in turn, evolving independentally along each lineage under a homogenous Poisson process. If the relationship between the trait states and the processes which they regulate were direct (e.g., the trait states were continuous variables that served as the rates for the speciation, extinction and dispersal processes), then we would describe our model as a compound Poisson model with the rate of change for the trait states as hyperparameter(s). However, in our formulation, the trait states are categorical variables, and their relationship to the rates of speciation, dispersal, and extinction are given by arbitrary user-defined functions that are part of our model. Our model can be understood as a modification or extension of the Dispersal-Extinction-Cladogenesis model (Ree et al. 2005; Ree and Smith 2008; Matzke 2014), by (a) the incorporation of the branching process that generates a phylogeny as part of the model (instead of treating the phylogeny as a parameter of the model supplied by the user); (b) the addition of the modeling of evolution of one or more traits under separate Poisson processes on the growing phylogeny; and (c) the addition of a set of rate functions that relate a given lineage’s trait states to the speciation, extinction and dispersal rates of that lineage.

Our model consists of the following components (described in detail following the summaries):

1. Geography: a set of areas and connections between them which define the spatial configuration of the system.
2. Traits: which defines the ecology or biology of the lineages insofar as they inform other aspects of the system (in particular, the speciation, extinction, and dispersal processes).
3. Lineages, representing an evolutionary operational taxonomic unit, corresponding to a node on a phylogeny.
4. A collection of lineage rate functions: a set of functions that provide rates or weights for the independent processes of speciation, extinction and range evolution of each lineage.
5. The process of anagenetic range evolution, which is actually the superposition of two independent processes: the process of *anagenetic range gain*, where a lineage adds an additional area to its range through dispersal and colonization of a new area; and the process of *anagenetic range loss*, where a lineage loses an area in its range through local extirpation. The latter also includes the generation of extinction events in the branching process of the phylogeny, i.e., corresponding to a death event in a birth-death model, as a special case, as a lineage is considered to go extinct once its range is reduced to the empty set.
6. The process of cladogenesis, which generates speciation events in the branching process of the phylogeny, corresponding to “birth” events in a classical birth-death model, where the phylogeny gains a lineage by an existing lineage splitting into two daughter lineages. This includes the generation of cladogenetic range evolution events, as each daughter lineage acquires a range based on various different kinds of speciation modes.
7. Termination conditions: which determines how long the process is run before the phylogeny is sampled.

#### Geography

The geography of our model can be represented as a directed graph, 𝒢, where vertices represent *areas* and arcs represent differential dispersal weights between areas. An “area” is a operationally-defined biologically-meaningful distinct geographical subunit of the study region: islands, mountains, zones of suitable habitat in a continuous landscape, etc. We distinguish between two classes of areas: focal and supplementary. Focal areas are geographically-demarcated units within the study region from which lineages will be sampled, while supplementary areas areas areas external to the study region which nonetheless share or contribute to the evolutionary history of the lineages within the study region. For example, in the study of an adapative radiation of a group of birds in an island system, the areas representing individual islands in the archipelago would be focal areas, while the continental source with which fauna is interchanged with the islands would be a supplemental area. The distinction between focal and supplemental areas is useful, as it allows for a better approximation of many island biogeographical systems, where the lineages evolving under quite different ecologies in distant regions contribute to the diversity and dynamics of the study region following introduction by stochastic dispersal. The relative diversity of each of the areas is a model parameter, representing the relative probability of which of the areas within a particular lineage’s range becomes the site of speciation in the event that the lineage splits. This allows for modeling of unequal diversity across areas, either as a reflection of area size or other factors. We use the notation *A*_*i*_ to refer to an the *i*-th area of the model.

The relative connectivity of the areas to each other are represented by the arcs of 𝒢. An arc is a directed edge connecting two vertices, going from a source or tail vertex to a destination or head vertex. In the geographical submodel, an arc connecting area *u* to area *v*, ***α***_〈*A_u_⇒A_v_*〉_ represents the weighting of a dispersal event from area *u* to *v*, relative to all other area-pairwise dispersal events. If all areas are equally connected, then all arcs will have the same weight, which normalizes to 1: ***α***_〈*A_u_⇒A_v_*〉_ = 1, *∀u v*. If some areas are not accessible from other areas, then the outgoing arcs of the latter toward the former will have a weight of 0. These weights will be multiplied by the lineage-specific dispersal weight of each lineage and the global disperal rate, ***δ***, to obtain the effective rate of dispersal of that lineage from the source area to the destination area. The relative weightings of arcs connecting areas allow for modeling of various geographical geometries, such as stepping-stone or equal-island configurations, as well as for more nuanced modeling of the dispersal of lineages from one (or more) continental sources to the areas that are the focus of the study.

#### Traits

Traits are characters associated with lineages. In the current discussion, we consider traits to represent discrete characters, though there is nothing in principle that prevents them from representing continuous characters. An arbitrary number of trait *types* can be modeled, with an arbitrary number of trait states per trait type. While traits can be used to represent any aspect of lineages, in this discussion we find it useful to consider them as representative of lineage ecology, just as areas represent the lineage geography.

Each lineage in the model shares the same suite of trait types, though, of course, the states of each particular trait type will vary from lineage to lineage. Each trait type evolves under its own Poisson process, and thus has its own distinct Markovian transition kernal, with the state of each trait type for a lineage evolving independently on each lineage under the Markovian transition kernel for that trait type. Daughter lineages inherit the trait state set of their parents, unmodified. This is, in fact, a fairly traditional model of phylogenetic character evolution (see, for example, Felsenstein 2004). We denote the set of all trait types defined in the model as **T**, and an individual trait type will be referenced by its labeling, as 𝒯_“label”_; for example, the “habitat” trait will be referred to as 𝒯_habitat_. We denote the the state space of an arbitrary trait with label “*t*”, 𝒯_*t*_, as ***σ***_t_, and the size of this state space as |***σ***_t_|. We denote as **Q**_t_ the *n × n* normalized instantaneous rate of change matrix for trait with label “*t*”, 𝒯_*t*_, where *n* is the size of the state space for trait type 𝒯_*t*_, *n* = |***σ***_t_|. In our model parameterization, the transition matrix **Q**_t_ is normalized so that the mean flux is 1, and describes the relative weights of state transitions, but to obtain the actual rate of transition this matrix is multiplied by a global trait transition rate. Each trait type has its own independent trait transition rate, and we denote the global trait transition rate for an arbitrary trait with label “*t*”, 𝒯_*t*_, as *q*_t_.

#### Lineages

A “lineage” in our system is an independent evolutionary unit, representing an operational taxonomic unit corresponding to an node on a phylogeny. Each lineage in our model can be characterized by two attributes: its “ecology”, which is a vector of trait states under the trait submodel (described below), and its “range”, which is the set of distinct areas (i.e., vertices in the geographical submodel, 𝒢, described below) in which it occurs. We use the notation *s*_*i*_ to denote the *i*-th lineage of the system, ***ξ***_t_(*s*_*i*_) to denote the current state of the trait with label “*t*” for lineage *s*_*i*_, and **R**(*s*_*i*_) to denote the set of areas currently in the range of lineage *s*_*i*_.

The traits of a lineage changes following a Poisson process as described above, with the states of each trait evolving independently on each lineage under the Markovian transition kernel specified for that particular trait. The range of a lineage changes through gain or loss of areas under two distinct process: anagenetic range evolution and cladogenetic range evolution. These are discussed in detail below.

#### Lineage Process Rate Functions

Our model requires that three functions be defined that regulate the processes of speciation, extinction, and range evolution for each lineage:

1. A *lineage birth rate* function, **Φ***_β_*(*·*).
2. A *lineage extirpation rate* function, **Φ**_*ϵ*_(*·*).
3. A *lineage dispersal weight* function, **Φ***_δ_*(*·*).

Each of these functions takes a particular lineage as an argument, and maps to a real value giving the weight or rate of the given process for that particular lineage at that particular point in time, based on evaluation of the properties of the lineage or some other criteria. Each of these functions can be fixed to a constant value irrespective of the lineage argument to implement a model where the given process is independent of any trait or property of the lineage. For example, we can specify that the birth rate of is a fixed value of 0.01 regardless of the lineage or any other criteria by specifying:

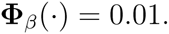

Alternatively, the functions can be more complex, and dependent on one or more trait states of the lineage. For example, if ***ξ***_t_(*s*_*i*_) returns the state value for trait “*t*” of lineage *i*, then we can construct a lineage dispersal weight function that evaluates to 0.5 if the “color” trait of lineage *i* has state “0”, and 1.0 if it has state “1”:

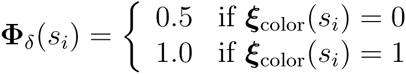

The functions can also reference other attributes of the lineage such as, for example, the number of areas in their range, or even the identities of particular areas. In this paper, however, we restrict our discussion to functions that only consider the traits associated with each lineage.

#### Anagenetic Range Evolution

Anagentic range evolution is when the range of a lineage changes, either by gaining an area or losing an area, during the lifetime of the lineage. The range of a lineage can expand anagenetically by adding an area through the process of colonization by dispersal. The global dispersal rate model parameter, ***δ***, determines the base probability of range gain through dispersal across the system. This model parameter is modified by the lineage-specific dispersal weight, as returned by the lineage dispersal weight function, and the weight of the connection arcs between the source and destination areas. The rate that a lineage *s*_*i*_ disperses from one particular area in its range, *A*_*u*_, *A*_*u*_ ∈ **R**(*s*_*i*_), to another particular area not already in its range, *A*_*v*_, *A*_*v*_ ∉ **R**(*s*_*i*_), is given by the product of the lineage dispersal weight, **Φ***_δ_*(*i*), the weight of the arc from *A*_*u*_ to *A*_*v*_, ***α***_〈*A_u_⇒A_v_*〉_, and the global dispersal rate, ***δ***:

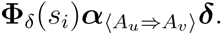

The *total* rate that a lineage *s*_*i*_ gains a new area not already in its range, *A*_*v*_, *A*_*v*_ ∉ / **R**(*s*_*i*_), is given by the sum of the rates of it dispersing from every one of the areas already in its range to the new area *A*_*v*_:

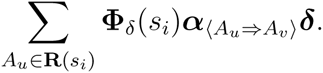

Conversely, the range of a lineage can contract anagenetically by losing an area through the process of extirpation. The probability that a lineage *s*_*i*_ loses an area already in its range is given by the lineage extirpation rate function applied to the lineage, **Φ***_ϵ_* (*s*_*i*_). If the lineage loses all areas in its range, i.e., its range is equal to the empty set, **R**(*s*_*i*_) = Ø, this means that it has gone globally extinct.

#### Speciation and Cladogenetic Range Evolution

The branching process that generates the phylogeny is driven by a birth-death process.

The lineage-specific birth rate, i.e. the rate that a particular lineage splits into two daughter lineages, is given by the lineage birth rate function **Φ***_β_*(·). The ranges of the daughter lineages are constructed based on one of the following speciation modes:

1. *single-area sympatric speciation* – both daughters inherit the entire ranges of their parent.
2. *sympatric subset speciation* – one daughter inherits the complete range of the parent lineage, while the other inherits a single area in the ancestral range
3. *peripheral allopatry* – one daughter inherits a single area with the parent’s range, while the other inherits all remaining areas
4. *multi-area vicariant speciation* – the ancestral range is partitioned unequally between the two daughter species
5. *founder-event “jump” dispersal* – one daughter inherits the entire range of the parent, while the other has its range set to a single a new area not in the parent range

These speciation modes correspond to modes defined in the BioGeoBears dispersal model (Matzke 2014), which extends the DEC model (Ree and Smith 2008) by adding a founder-event “jump” dispersal. At every cladogenesis event, one of these speciation modes is selected under a model-specified probability distribution to determine the ranges of the daughter lineages. In all work reported here, we placed a probability of 0 on the founder-event “jump” dispersal speciation mode, and equal probability on all the other modes. This restriction is so that in all the analyses we report on here, the dispersal sub-model exactly replicates the DEC (Ree et al. 2005; Ree and Smith 2008) model, without the BioGeoBears (Matzke 2014) extension.

If there is a need to model different levels of the diversity in the various areas, different diversity weights can be given to each of the areas. These diversity weights determine the probability that a particular area becomes the site of a new species in speciation modes 3, 4, and 5, for example. In all the work reported here, we set the diversity levels of all areas to be equal.

The lineage-specific extirpation rate, **Φ**_*ϵ*_(·), gives the rate by which a lineage loses areas from its range. If the range of a lineage is reduced to the empty set, then it is considered to have gone extinct. Thus, while there is real extinction in our model in addition to speciation, the death rate parameter is not an explicit primary parameter of our model as it is in the standard birth-death model. If there is a need to establish full correspondence between the extirpation rate as given in our model and the death rate standard birth death-model (for, e.g., calibration purposes), the lineage extirpation rate function can be modified to return the global extinction death rate multiplied by the number of areas currently occupied by the lineage, as the global extinction probability is the joint probability of the lineage being exirpated from every area in its range simultaneously.

#### Termination Conditions

The phylogeny generated by this model is a dynamic entity, with every aspect changing over time: size (number of leaves/extant lineages), branch lengths, shape/structure, as well as the distribution of the lineages in both geographical as well as trait space. At the termination of the phylogeny generation process, the state of the phylogeny is sampled and stored as the final output of the process. There are three ways to determine the duration for which the process runs before termination:

1. The process can be run for a fixed length of time.
2. The process can be run until a specified number of tips (extant lineages) are generated in the focal areas (i.e., the number of leaves that include at least one of the focal areas in their ranges).
3. As above, but under the General Sampling Approach of Hartmann et al. (2010), in which the process is run until a sufficiently large number of lineages are generated in the focal areas, and then a “snapshot” of the phylogeny at a time when the number of lineages in the focal areas are equal to the target number of lineages are selected with uniform probability.

### Simulations under the “Archipelago” Model

#### Initialization

At simulation time, *t* = 0, we initialize the system with a seed lineage. This seed lineage constitutes the beginnings of our phylogeny, and it has its (initial) edge length set to 0, and is added to the set of extant lineages, **L***^*^*.

For each of the traits being simulated, the initial trait state of the seed lineage is selected at random from the stationary distribution of the trait. In all the work reported here, we use a simple equal-rates model for the evolution of all the traits, so the initial trait states of each of the seed lineages in our simulations were sampled with uniform random probability.

We also initialize the range of the seed linage with a single area. While we could in principle select an area at random (perhaps in proportion to the diversity weight of the area), in all work reported here, we used the first area defined as the initial area of the seed lineage, as the first area so defined corresponded to the supplemental continental area being modeled as an external source.

#### Life-Cycle

Each of the extant (current or tip) lineages in “Archipelago” evolves independently of all other extant lineages under its own complex of superpositioned independent speciation, extirpation, dispersal, and trait evolution processes. The phylogeny as a whole, in turn, evolves under the superposition of each of these superpositioned processes across all the extant lineages. At any point in time of the simulation, the sum of the rates of each possible event that may occur under each of the independent processes across all extant lineages is the rate of *any* single one of these events occurring. The rate of each of these events normalized by the sum of these rate, in turn, gives the probability of that particular event occurring conditional on *any* (single) event occurring.

Consider a lineage, *s*_*i*_, in the set of extant lineages, *s*_*i*_ ∈ **L***^*^*. This lineage may:

1. Split into two daughter lineages at the lineage-specific birth rate of **Φ***_β_*(*s*_*i*_).
2. Lose an area from its range at the lineage-specific extirpation rate of **Φ**_*ϵ*_(*s*_*i*_).
3. Gain an area in its range at a rate of:

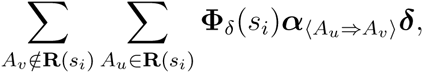

where **R**(*s*_*i*_) is the set of areas in the range for lineage *s*_*i*_, **Φ***_δ_*(*s*_*i*_) is the lineage-specific dispersal weight for lineage *s*_*i*_, ***α****_〈_u_⇒A_v_〉_* is the weight of the arc connecting *A*_*u*_ and *A*_*v*_, and ***δ*** is the global dispersal parameter.
4. Evolve one of its traits, 𝒯_*i*_, 𝒯_*i*_ ∈ 𝒯, at a rate of:

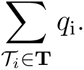

The sum of the rates of the above events across all extant lineages in **L***^*^* gives the rate of *any one* of these events occurring in any extant lineage across the phylogeny, ***λ^*^***:

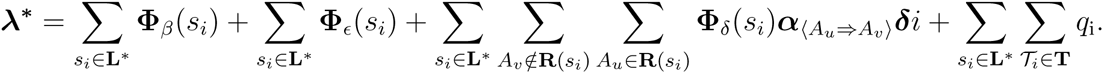

The waiting time, *w*, until any of the above enumerated events occurs in the phylogeny is sampled from an exponential distribution with a rate given by ***λ^*^***:

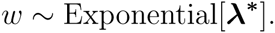

If the termination condition is set to terminate at fixed time, and if the current simulation time plus the waiting time exceeds this, then the difference between the the fixed termination time and the current simulation time is added to all extant lineages in **L***^*^*, and the simulation is terminated with current phylogeny as the final phylogeny. Otherwise, this waiting time is added to the edge lengths of all extant lineages in **L***^*^* as well as the current simulation time, *t*. If the current number of extant lineages in the focal areas is equal to the number of target taxa specified in the termination criteria under the General Sampling Approach of Hartmann et al. (2010), then a snapshot of the phylogeny at the current time is stored, along with the waiting time as the duration associated with it.

Following this, an individual event is selected to be realized with probability given by the rate of the individual event normalized by the rate of any event, ***λ^*^***. For example, the probability that a particular extant lineage, *s*_*i*_, *s*_*i*_∈ **L***^*^*, speciates by splitting into two daughter lineages, conditional on an event having taking place is given by:

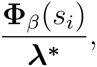

while the probability that it gains a new area not already in its range, *A*_*v*_, *A*_*v*_ ∉ **R**(*s*_*i*_), is given by:

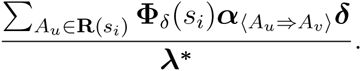

If the event is a speciation event for a particular lineage, *s*_*i*_, then two new lineages are created with their edge lengths initialized to 0.0 and their parent set to *s*_*i*_, and added to the set of extant lineages, **L***^*^* children of *s*_*i*_, while *s*_*i*_ itself is removed from that set. The new daughter lineages inherit the entire set of traits from *s*_*i*_, while their ranges are set according to the speciation mode as described above.

If the event is an extirpation event for a particular lineage, *s*_*i*_, then an area is selected at random to be removed from the set of areas in its range, **R**(*s*_*i*_). If this results in its range being reduced to the empty set, **R**(*s*_*i*_) = *Ø*, then the lineage is considered to have gone extinct and is removed from the set of extant lineages, **L***^*^*.

If the event is a range gain event, where a particular lineage, *s*_*i*_, gains a new area, *A*_*z*_, then that area is simply added to the set of areas in the range of the lineage, **R**(*s*_*i*_). If the event is a trait transition, on the other hand, than the given trait state for the lineage is appropriately adjusted.

Once the selected event has been executed and processed, the number of extant lineages in the focal areas are counted, i.e., the number of lineages that include at least one focal area in their range. If the termination condition is set to a fixed number of lineages in the focal areas, and this current number of extant lineages in the focal area does indeed equal or exceed this, then the simulation is terminated with the current phylogeny as the final phylogeny. If the current number of extant lineages equals the number of maximum number of lineages specified in the termination criteria under the General Sampling Approach (Hartmann et al. 2010), then one of the stored phylogeny snapshots is selected with probability proportional to its duration, and the simulation is terminated with this as the final phylogeny.

#### Termination and Completion

The final phylogeny of simulation is pruned to exclude any lineages that do not occur in the focal areas, and this is saved as the ultimate product of simulation. This phylogeny will be an ultrametric phylogeny, with the edge lengths reflecting the duration of the respective lineages. Each of the tip lineages will be labeled to indicate the states of each trait associated with it as well as its range at the time it is was sampled.

### Model Selection by Simulation-Trained Discriminant Analysis of Principal Components (DAPC) Classification

Carrying out full-likelihood inference under this model would be challenging, both for theoretical as well as practical reasons. Depending on the various lineage-specific rate and weight functions, the rates of the various processes can vary across both lineages as well as time in a stochastic and unpredictable way. Here we present a simulation-based likelihood-free model choice procedure as an alternative to full-likelihood Bayesian or frequentist approaches, where we use a multivariate discriminant analysis function to classify the target or empirical data with respect to one of a number of competing generating models.

Our procedure consists of the following steps:

1. Simulation of data (phylogenies) under regimes corresponding to hypotheses in the competing model set, with nuisance parameters estimated from target datum that we wish to classify.
2. Calculation of summary statistics on simulated data to obtain a data set that can be used to train a classification function.
3. Construction of a discriminant analysis function based on principal components extracted from the training data set.
4. Assessment of the performance of the summary statistics by inspection of posterior prediction of the training data set.
5. Application of the discriminant analysis function to the original data to classify it with respect to the competing models.

#### Simulation of Training Data

We simulate data under the “Archipelago” model using the Python (Van Rossum and Et Al. 2010) package “`archipelago`” (Sukumaran 2015, formal citation in prep.). Data is simulated under different classes of parameter regimes, with each class of regimes corresponding to a particular evolutionary hypothesis in our candidate set of models. As each datum n the data set is thus generated under known conditions, the entire collection serve as a training data set for the construction of the discriminant analysis classification function which will be used to classify the target or empirical data.

Consider, for example, determining whether or not the habitat-association of lineages influences their dispersal capacity. We would define two classes of dispersal regimes: a habitat-unconstrained dispersal vs. a habitat-constrained dispersal. Under the habitat-unconstrained regime, all lineages disperse between areas at the same rate, regardless of their ecology in terms of habitat-association. Under the habitat-constrained reigme, only lineages associated with a particular habitat type are allowed to disperse. Figure 1 illustrates these two regimes applied to a two-island system, where only lineages with associated with an open or disturbed habitat ecology are capable of dispersing, while lineages associated with interior habitats (e.g. undisturbed, primary forest) cannot.

**Figure 1.**
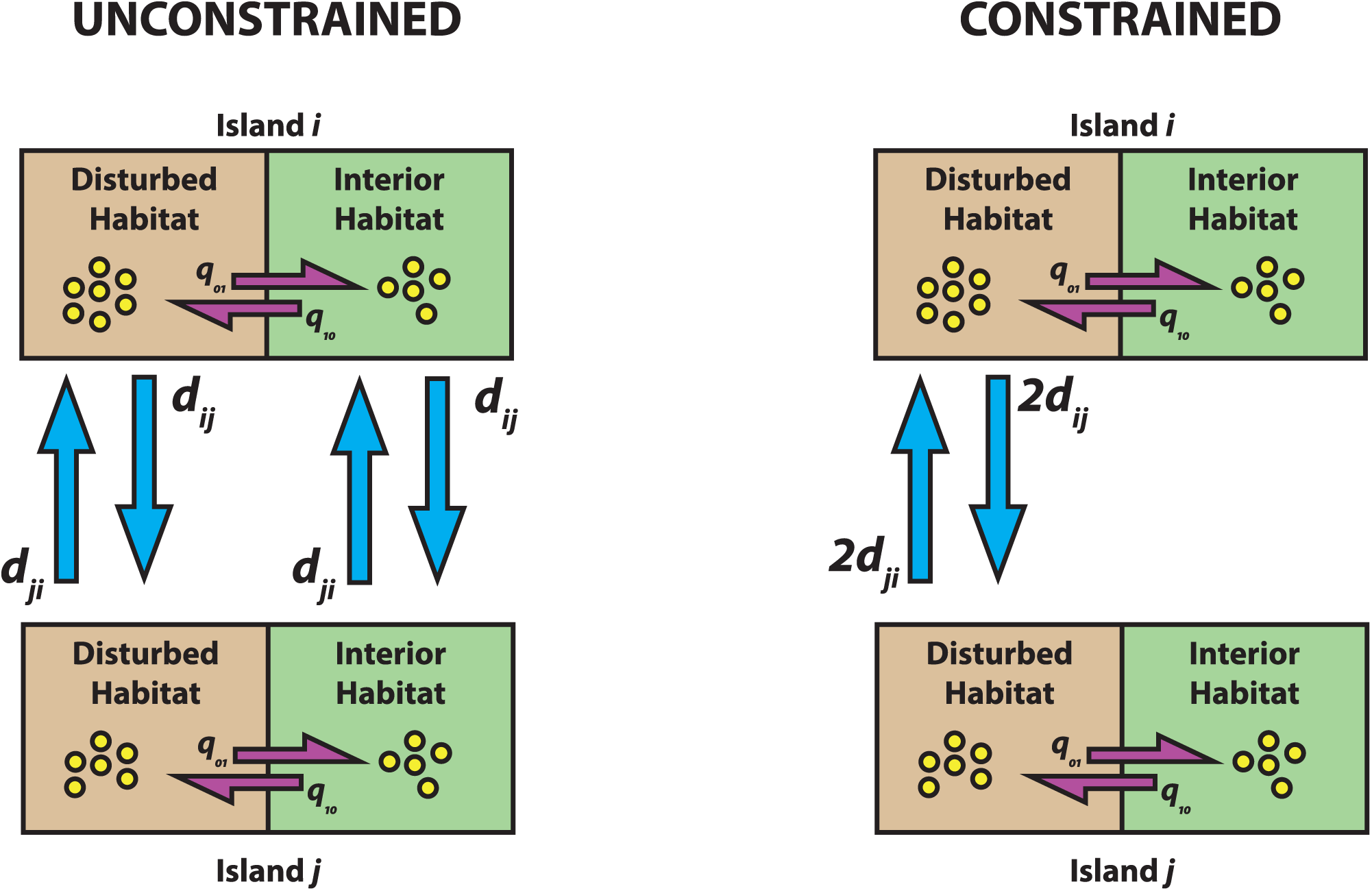
Schematic for study design to test if dispersal is constrained by habitat. In the “unconstrained” regime (left), lineages are free to disperse between islands regardless of their habitat trait. In the “constrained” regime (right), only lineages associated with the interior (undisturbed) habitat can disperse. Lineages evolve from the constrained habitat to the unconstrained under the same rate, *q*_10_ = *q*_01_. Dispersal rates in the constrained regime are weighted twice the dispersal rates in the interior regime to ensure that, on the average, the total flux between the constrained and unconstrained regimes are equal. This two-island system can be increased to an arbitrary number of islands.

Under the “Archipelago” model, we would define a trait, “habitat”, 𝒯_*habitat*_ that had two states, “open” and “interior”, ***σ***_habitat_ = {open, interior}. In all the studies presented here, we assume equally-weighted transition rates between all trait states in all our trait evolution models, such the rate of transitions from the “open” trait state to the “interior” is equal to the rate of transitions from the “interior” to the “open” state.

In the case of the habitat-unconstrained dispersal regime, we would define a fixed-value lineage dispersal rate function,

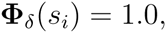

which would mean that every lineage would disperse equally under rates given by the product of the global dispersal rate, ***δ***, and the relevant dispersal weights between areas, regardless of whether the habitat trait associated with the lineage was “open” or “interior”.

On the other hand, in the case the habitat-constrained dispersal regime we would define a lineage dispersal rate function that only allows dispersal if the lineage’s habitat trait was “open”:

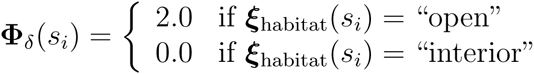

Here, the effective dispersal rate for any lineage with the “open” habitat trait is 0.0, so that these lineages will not gain any areas due anagenetically, regardless of the connections between areas or the global dispersal rate, ***δ***. Only lineages with the “open” habitat trait will gain areas in their range anagenetically, at an effective rate of 2.0***δ*** multiplied by the dispersal weights of all the areas in which they occur. Note that the dispersing lineages in the constrained model have their rates of dispersal weighted twice as much as the lineages in the unconstrained model, to ensure that the average dispersal over both constrained and unconstrained dispersal regimes are equal. We use a factor of 2 because, under a trait-transition model in which transition rates between all states are equal, we expect an equal number of lineages to be found associated with each of the two habitat states, and thus in the constrained model, on the average only half the lineages are dispersing in relation to the unconstrained model. This means that for the total dispersal flux to remain equal when estimated from data generated under both systems, we have to weight the constrained regime dispersal rates to compensate for the fact that only half the lineages are dispersing. We show later that this weighting does, indeed, result in correct behavior, as measured when dispersal rates are re-estimated from the simulated data under both regimes.

In all other respects, apart from the lineage dispersal weight function, the models of unconstrained and constrained dispersal regimes are identical: they are calibrated to share the same (speciation, extinction, dispersal, and trait evolution) process parameters, number of focal areas, supplemental areas, as well as termination conditions. Thus in *both* the unconstrained and constrained dispersal regime models, we set the lineage specific birth rate and extirpation rate functions to return fixed values, 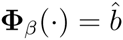 and 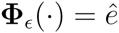, where 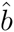 and 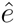 are the birth rate and local extirpation rate are estimated from the empirical data, using DendroPy (Sukumaran and Holder 2010) and BioGeoBears (Matzke 2014), respectively. Similarly, the global dispersal rate, ***δ*** for both the unconstrained and constrained dispersal regimes is set to the same value, 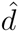, which is also estimated from the empirical data using the “`R`” package “`BioGeoBears`” (Matzke 2014), as is the trait transition rate, 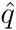, using the “`R`” package “`GEIGER`” (Harmon et al. 2008). The landscape for both dispersal regimes, i.e., the number of areas would reflect the number of areas in the study system (e.g., islands in an archipelago, or archipelagos of a region, depending on the scale of the study), with optional supplementary areas (e.g., adjacent continent) if required. The termination condition can be constructed in number of ways supported by the model, and for all the analyses reported here, we choose to terminate when the number of extant lineages in the focal area equals the number of tips in the observed or target phylogeny.

With simulate a 100 replicated under each of the regimes, habitat-unconstrained and habitat-constrained dispersal. Each simulation yields a phylogeny containing information not only on speciation times, but on ranges (incidence in terms of islands or areas) and habitat associations of each lineage.

#### Calculation of Summary Statistics on Training Data

The simulations described in the previous step produced a set of phylogenies under each of the model regimes. For each of the simulated phylogenies, we calculate a vector of summary statistics. The actual choice of summary statistics would, of course, vary based on application. The principal criteria for summary statistic selection here is that they capture some of the patterns generated by the processes that we are studying. In our motivating examples, as we are interested in the relationship between a particular trait (habitat-association) and geographic distribution, and we use statistics based on the Mean Phylogenetic Distance (MPD) and Mean Nearest Taxon Distance (MNTD) concepts from community ecology (Webb 2000). We calculate two variants of each of these statistics, depending on whether we define a “community” based on area (i.e. island) or habitat (i.e., open vs. interior habitats). For the area-based community scores, we calculate the mean and discard the individual area-based community scores; this is because in the analyses discussed here, we consider the areas/islands as exchangeable as a simplifying assumption, and more complex analyses may gain form retaining the individual island scores. For the habitat-based community scores, we retain the ones defined on considering each individual habitat as a community, as well as the mean of these scores. Furthermore, for each of these statistics, we calculate both weighted as well as unweighted distances, with edge lengths being considered when calculating distances between taxa in the former and just the number of edges between taxa being used in the latter. All weighted scores are normalized with respect to total tree length, while all unweighted scores are normalized with respect to the total number of edges on tree. In addition to the actual MPD or MNTD Z-score, we also use the p-value and variance of the scores, with p-value calculated using a null-distribution obtained from shuffling the taxon labels. The actual number of summary statistics with this scheme will, of course, vary depending on the number of trait states (i.e., habitat category types). For example, with two habitat categories as trait states, open and interior, we obtain a total of 47 individual summary statistics. All these community-ecology summary statistics were calculated using the “`R`” package “`picante`” (Kembel et al. 2010). The package “`archipelago`” (Sukumaran 2015) provides a convenient script that automatically calculates the full suite of summary statistics given a phylogeny or set of phylogenies. A detailed list of summary statistics used in this study will be made available in the supplemental materials.

#### Construction, Assessment, and Application of the DAPC Function

We construct a discriminant analysis function using the simulated data as a training data set, using the dispersal model regime as the grouping factor. We do not construct this function based on the summary statistics of the training data set directly, but rather on principal components extracted from the summary statistics, i.e, a Discriminant Analysis of Principal Components, or DAPC, approach (Jombart et al. 2010). The discriminant analysis function that will be used to classify data under one of the model regimes is constructed using the “`dapc`” method in the “`R`” package “`adgenet`” (Jombart 2008), which is wrapped in a convenient script provided by the “`archipelago`” (Sukumaran 2015) package.

The number of principal component axes retained or used by the discriminant analysis function is an analysis design choice. Too many principal components will result in over-fitting to the training data set, while too few will result in lack of power (Jombart et al. 2010). Here we select the number of the principal component axes to be retained for the discriminant analysis function using a heuristic optimization criteria, where, we attempt to (re-classify) the training data set elements using discriminant functions constructed on a different numbers of principal components retained, and select the number of principal components to retain that maximizes the proportion of correct (re-)classification of the original training data elements. The “`archipelago`” (Sukumaran 2015) package script that carries out the classification procedure also carries out this heuristic optimizaiton of the number of principal components to retain by default.

The performance of the DAPC function can be assessed in terms of its success rate in re-classifying the training data: the discriminant analysis function is applied to each of the vectors of summary statistics to estimate the posterior probability of being generated under each of the competing model regimes. This performance can be measured in two ways: the mean posterior probability the true model regime, as well as the proportion of instances in which the true model was the preferred model regime.

The application of the DAPC function to classify the empirical data is carried out with the “`predict.dapc`” method of the “`adgenet`” (Jombart 2008), which returns the posterior probability of the input data belonging to each of the groups (i.e., candidate model regimes) defined in the training data set used to construct the DAPC function, as well as a classification of the input data into one of the groups based on the highest posterior probability. This classification essentially gives as the model chosen by our inference method based on the empirical data and the training data set.

The entire process of ingesting a training data set, selecting the number of PC axes to retain, constructing a DAPC function, and applying it to an empirical data set to yield the model choice results is wrapped up by an application script in the package “`archipelago`” (Sukumaran 2015).

### Assessment and Validation of Model Choice Performance

We assessed the performance of our model-selection by simulation-trained discriminant analysis classification by simulating independent training and test data sets, and evaluating the proportion of correct classifications (model selection) when the independent test data sets were analyzed using DAPC functions constructed on the trained data sets.

We used the a single two-state trait system, where our hypothetical trait, “habitat” could be either “open” or “interior”. We attempted to distinguish between two competing dispersal regime models, “unconstrained” vs. “constrained”, across a broad range of parameter space. In the “unconstrained” regime model, lineages could disperse between areas regardless of their “habitat” trait state, while in the “constrained” dispersal regime model, lineages could only disperse between areas if their “habitat” trait was “open”. We used a landscape consisting of four focal areas and one supplemental area, with simulations running until 200 lineage were generated in the focal areas. The weighting of the dispersal rates between all areas were set to be equal, as was the diversity weights.

We explored the performance of our method over a range of process parameter space. We defined our parameter space exploration in terms of birth rates, i.e., we explored suites of parameters all scaled to a particular birth rate. The birth rates we used are given by the set, **b**, where **b** = *{*0.0015, 0.0030, 0.0060, 0.0150, 0.0300, 0.0600*}*. For *each* birth rate, *b, b ∈* **b**, all distinct combinations of the following parameters were explored:

- extirpation rates, *e*: 0, 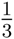*b*,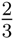*b*.
- dispersal rates, *d*: *b ×* 10^-1^, *b ×* 10^*-*0.75^, *b ×* 10^*-*0.5^, *b ×* 10^*-*2.5^, *b ×* 10^0^, *b ×* 10^2.5^, *b ×* 10^0.5^, *b ×* 10^0.75^, *b ×* 10^1^.
- trait transition rates, *d*: *b ×* 10^*-*1^, *b ×* 10^*-*0.75^, *b ×* 10^*-*0.5^, *b ×* 10^*-*2.5^, *b ×* 10^0^, *b ×* 10^2.5^, *b ×* 10^0.5^, *b ×* 10^0.75^, *b ×* 10^1^.

For each distinct combination of parameters above, we then simulated a 200 target data sets, with 100 target data sets under the “unconstrained” dispersal regime and 100 target data sets under the “constrained” dispersal regime.

Each of these target data sets were profiled with respect to their major process parameters. The maximum-likelihood estimates of the birth rate under a pure-birth model were computed using DendroPy (Sukumaran and Holder 2010), the maximum-likelihood estimates of the dispersal rates were computed under the DEC model using BioGeoBears (Matzke 2014), and the maximum-likelihood estimates of the habitat trait transition rate under a equal rates model were computed using GEIGER (Harmon et al. 2008). This was to establish that the (a) data that we were generating were actually under the process parameters, and (b) that our weighting of dispersing lineages in the constrained system correctly recovered the estimated dispersal rates.

For each target data set, we then generated a training data set of 200 replicates, with a 100 replicates under the “constrained” dispersal regime and the other 100 replciates under the “unconstrained” dispersal regime, using the *true* model parameters (i.e., not the estimated ones recovered during the profiling in the previous step). We acknowledge that using the estimated parameters would provide a more stringent assessment of the performance of the method, as our method then has to deal with estimation error in the calibration of the process parameters when simulating the training data sets. However, we suggest that our approach is justified because, as the results of the profiling described above show, our simulations produce data that result in estimates of process parameters that are very close to the true parameters. A discriminant analysis classification function was constructed on principal components calculated on each training data set, with the number of principal components retained selected by the optimality criteria described above. The proportion of correctly-classified test data sets was taken as the primary performance metric, but we also recorded the posterior probability of the true and false models to understand how these varied across parameter space.

### Sensitivity of Model Choice to Process Parameter Calibration Value Errors

In our approach as presented here, we “calibrate” the simulations that generate the training data sets to match the target data set with respect to the birth, dispersal, and trait evolution process parameters. In practice, when attempting to classify empirical data, these parameters are not known and must be estimated. We are interested in characterizing the sensitivity of our model choice approach to errors in these calibration parameters. To do this, using the same two-trait, four-focal area, and one-supplemental area system described previously, we simulate a target data set under a set of known process parameters, and then attempt to classify it with respect to the model regime using training data sets that deliberate mis-specify process parameters values. We use the following set of process parameters as baseline values for the process parameters: birth rate, *b* = 0.03; extirpation rate *e* = 0.00; global dispersal rate *d* = 0.03; and trait evolution transition rate *q* = 0.03. The training data set will use the same baseline values except one of the parameters will have an error introduced. We analyze the effect of erros in the birth rate, dispersal rate, and trait trait evolution transition rate separately. In each case, the target data set will be generated under the baseline values, and the training data set will be generated with the parameter being studied changed by a factor (and all other parameters fixed to the baseline values). The factors that we use result in parameter values that range from two orders of magnitude less than, to two orders of mangnitude greater than the true (baseline) values, in increments of quarter orders of magnitude: 10^-2^, 10^*-*1.75^, 10^*-*0.5^, 10^*-*0.25,^ 10^0^, 10^0.25^, 10^0.5^, 10^0.75^, 10^1^, 10^1.25^, 10^0.5^, 10^0.75^, 10^2^. We repeat each analysis 200 times, with a 100 replicates in which true model of the target data set was “unconstrained” and a 100 replicates in which the true model of the target data set was “constrained”. Performance was assessed by measuring how the posterior probabilities of the true and false models varied across these different magnitudes of errors in each of the parameters.

### Analysis of Trait-Dependent Dispersal in Island Bird Radiations

We applied our model and inference method to try and identify whether dispersal is a constrained by traits in radiations of birds in Wallacea (figure 2), focussing on two traits in particular: habitat type and trophic-level.

**Figure 2.**
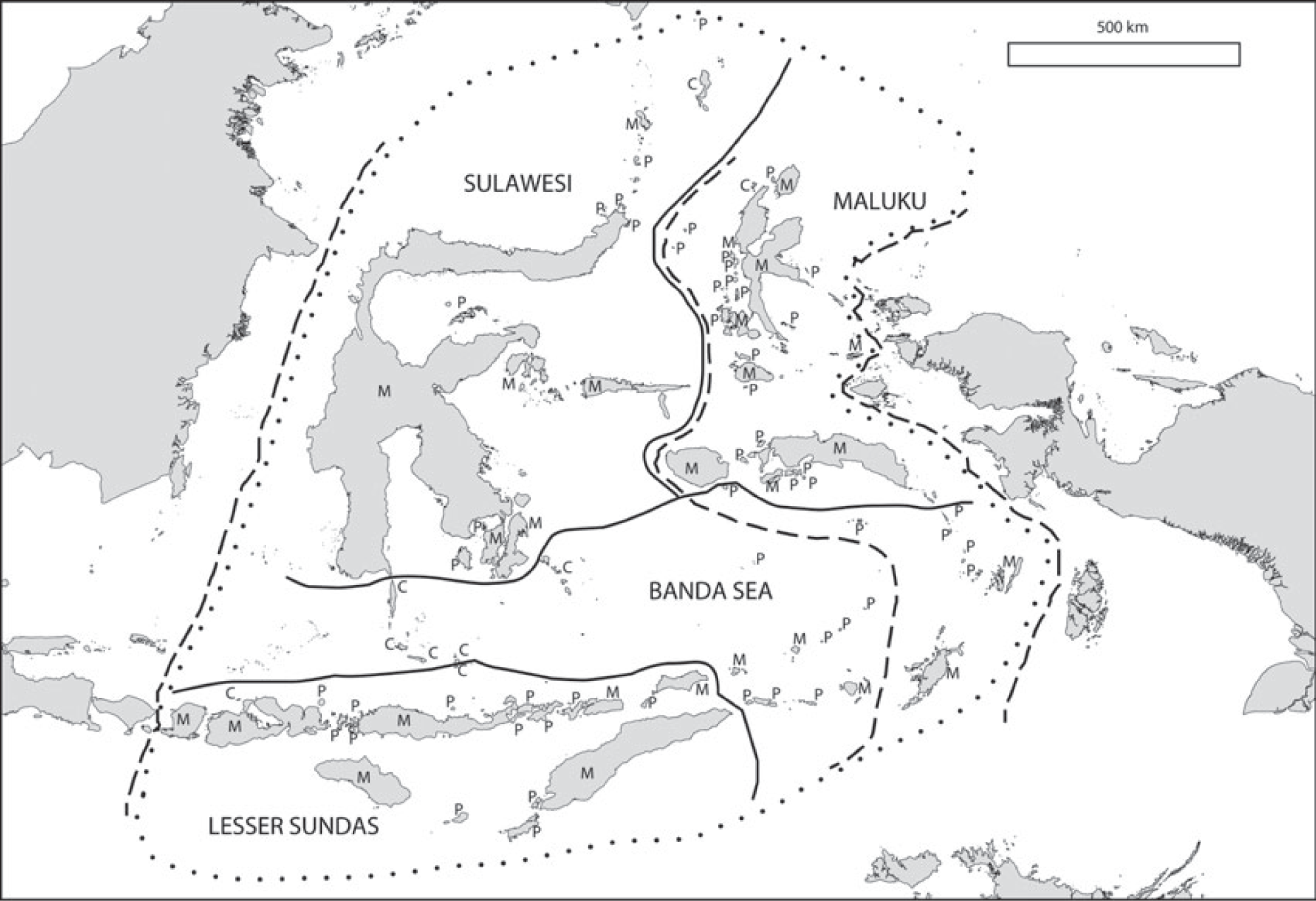
The biogeographical “modules” of Wallacea of Carstensen et al. (2012), with module names in small caps. The broken lines mark, from west Wallaces Line, Webers Line and Lydekkers Line. Figure from Carstensen et al. (2012). See Carstensen et al. (2012) and Carstensen et al. (2013) for more details.

#### Geography

Each trait was analyzed separately in a series of independent analysis using the same geographical submodel. The basic geographical units or “areas” of the analyses corresponded to the “modules” of Carstensen et al. (2012, 2013): Banda Sea, Lesser Sundas, Maluku, and Sulawesi (figure 2). These areas constituted the primary focal areas of the study, i.e., the areas from which lineages were be sampled. However, in addition, we added supplementary areas to the model to represent the Sundaland and Sahul continental sources which contributed to the evolutionary history, dynamics, and diversity of the region, but from which lineages were not be sampled for analysis. This more authentically approximated the sampling of the empirical data we were analyzing. In three independent sub-analyses for each trait, we used zero, one, and two supplementary areas respectively, followed by a fourth independent subanalysis for each trait in which the training data sets were the union of the previous three (i.e., essentially integrating over the number of supplementary areas with an equal prior on each).

#### Phylogeny

We downloaded 10000 post-burnin MCMC samples of the Jetz et al. (2012) global avian phylogeny using the Hackett backbone from [source], pruned to only include the bird species in the Wallacean fauna studied by Carstensen, that could be reliably mapped to the Jetz nomenclature. These trees were originally estimated using “`BEAST`” (Drummond and Rambaut 2007), and are ultrametric with an arbitrary calibration point of 100 time units at the root. A maximum clade credibility tree (MCT) was calculated on the MCMC trees using DendroPy/SumTrees (Sukumaran and Holder 2010). This resulting tree constituted the base operational phylogeny for this suite of analyses. For each trait-specific analyses set, the base operational phylogeny was modified by pruning species for which no data or no suitable data was available (see below for description of criteria, and Supplemental Materials for lists of species).

#### Habitat-Dependent Dispersal Analyses Trait Configuration

The original Carstensen study associated each species with one of six habitat categories:

- (H1) Interior: species only occupying interior forest
- (H2) Open-forest: species occupying interior forest and/or open forest
- (H3) Coastal: species only occupying littoral and/or open habitats
- (H4) Open: species only occupying open habitats
- (H5) Generalist: species occupying all four habitat types
- (H6) Other: species that did not fit into any of the previous five categories

In this study, we collapsed these habitats into two categories, “open” and “interior”, and model “habitat” as a two-state trait associated with, and evolving along, lineages. We categorized any species that occurs in open or disturbed habitat as a “open area” species, in the sense that they have access to open areas and thus are at least partially subject to dispersal regimes associated with open areas. On the other hand, only species that are strictly restricted to interior habitats are considered “interior area” species: they are excluded from any dispersal regimes associated with open areas. This means that we considered birds associated with habitats 2 through 5 as open area species, while birds associated with habitats 1 were categorized as interior area species. We discarded (and pruned from the phylogeny) any species associated with habitat category 6 (“Other”), as this was a polyphyletic category, and we are modeling the habitat-trait as an evolutionary character.

We define two dispersal regimes categories based on the lineage dispersal weight responses to the habitat trait. The first is the “unconstrained” dispersal regime model, where the lineage dispersal weight function is a fixed value of 1.0. The second is the “constrained” dispersal regime model, where the lineage dispersal weight function is conditional on the habitat trait state of the lineage: if the lineage habitat trait state is “open”, then it the dispersal weight is 2.0, but if it is “interior”, then the dispersal rate is set 0. This latter configuration thus only permits lineages with the “open” habitat trait to disperse.

#### Trophic-level Dependent Dispersal Trait Configuration

The original Carstensen study associated each species with one of nine guilds: frugivores, granivores, herbivores, nectarivores, insectivores, invertebrates, omnivores, piscivores, and carnivores. We modeled these guilds as trophic-levels using a two-state trait system, with the trait, “trophic level” being either “low” or “high”. The “low” trophic-level consisted of the “frugivore”, “‘granivore”, “herbivore”, “nectarivore”, “insectivore”, and “omnivore” categories, while the “high” trophic-level consisted of the rest. As before, we define two dispersal regimes categories based on the lineage dispersal weight responses to the trophic-level trait: the “unconstrained” dispersal regime model and the “constrained” dispersal regime model. In the “unconstrained” dispersal regime model, the lineage dispersal weight function is a fixed value of 1.0. In the “constrained” dispersal regime model, the lineage dispersal weight function is conditional on the trophic-level trait state of the lineage: if the lineage habitat trait state is “low”, then it the dispersal weight is 2.0, but if it is “high”, then the dispersal rate is set 0. This latter configuration thus only permits lineages with the “low” trophic-level trait to disperse. We pruned from the phylogeny any species for which no trophic level or guild information was available.

#### Simulation Calibration

The simulation calibration process parameters were then estimated separately on each trait-specific pruned phylogeny: the birth rate, *b*, under a pure-birth model using DendroPy (Sukumaran and Holder 2010); a trait evolution transtion rate, *q*_habitat_ or *q*_trophic-level_, depending on the trait being analysed, under an equal-rates model using the GEIGER package (Harmon et al. 2008); as well as a global dispersal rate, *d*, and an extirpation rate, *e*, under a DEC model (Ree et al. 2005; Ree and Smith 2008) using the “`BioGeoBears`” package (Matzke 2014). The global dispersal rate parameter of the “Archipelago” model was set to the estimated dispersal rate, ***δ*** = *d*, while the lineage birth and death rate functions were set to return the estimated birth rate and extinction rate values regardles of lineage, **Φ***_β_*(*·*) = *b* and **Φ**_*ϵ*_(*·*) = *e*.

#### Training Data Generation and Classification Analysis

For each of the analyses, we generated an independent training data set consisting of 100 replicates under the “unconstrained” dispersal regime and another 100 under the “constrained” dispersal regime, using “`archipelago`” (Sukumaran 2015). Each simulation was set to terminate when the number of lineages generated in the focal areas were equal to the number of lineages in the pruned trait-specific phylogeny. For each training data set, we constructed a discriminant analysis function classification function and applied it the empirical data to classify it with respect to the generating regime, “unconstrained” or “constrained”, following the approach described previously. We then conducted a fourth analyses for each trait, constructing a discriminant analysis classification function based on the union of the the previous three training data sets, and applying this to the empirical data as well. In all cases, the number of principal component axes retained was selected using the heuristic we describe above, i.e., maximizing the proportion of correctly-classified elements when the discriminant analysis function is re-applied to the training data set.

## RESULTS

### Process Parameter Estimates Under Unconstrained and Constrained Dispersal Regimes

Figure 3a compares the birth rates under which the target data was simulated to the maximum-likelihood estimates of the birth rate under a pure-birth model, grouped by dispersal regime. In general, the results are satisfactory: the estimates of the birth rate track the simulation birth rate closely, and there are no strong differences between the dispersal regimes. The long tails of the violin plots are indicative that most parts of the parameter space explored included moderate to very strong extirpation process which results in extinctions that are not accounted for by the pure-birth model.

**Figure 3.**
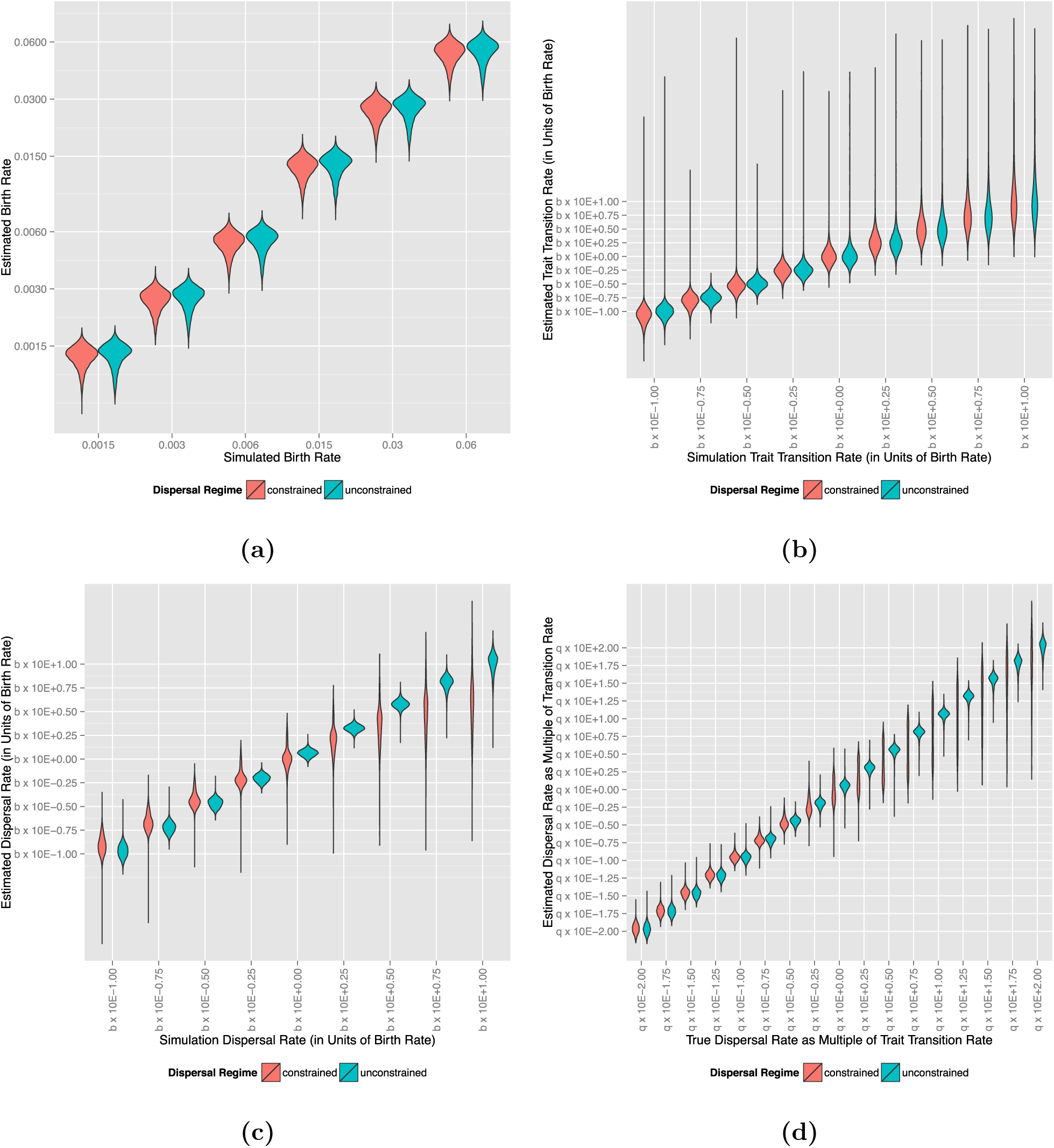
Maximum-likelihood estimates of various process parameter values compared to true (simulation) values to verify correct behavior of simulations, showing that the “archipelago” simulations are generating data as expected with respect to the (a) diversification, (b) trait evolution, and (c) dispersal parameters. In all cases, estimated parameter values generally track the simulation parameter values in both the “unconstrained” and “constrained” regimes. In the case of the “dispersal” parameter, however, the variance is greatly increased in the “constrained” regime, and this is related to variance in the trait transition rate, and thus stochasticity in the number of dispersing vs. non-dispersing lineages.

Figure 3b compares the habitat trait transition rates under which the target data was simulated to the maximum-likelihood estimates of the trait transition rates under an equal-rates model, grouped by dispersal regime. Note that we are expressing the transition rates in units of log of the (true) birth rate. As with the birth rates, the estimated trait transition rates generally track the true trait transition rate closely, with no strong differences between the dispersal regimes. As the true trait transition rate increases, however, to an order of magnitude higher than the birth rate, the variance in the estimates trait transition rates increases dramatically, with a very strong over-estimation bias.

Figure 3c compares the dispersal rates under which the target data was simulated to the maximum-likelihood estimates of the dispersal under a DEC model, grouped by dispersal regime. As in the previous, we express the rates in units of log of the (true) birth rate. Again, the estimated rates *generally* track the true rates, and the centers of mass of the estimates under both regimes are generally co-located except in the most extreme dispersal rates (i.e., with dispersal rates half or a full order of magnitude higher than the birth rates). However, the differences in variances between the dispersal regimes are striking. In particular, the variance in the estimates of the dispersal rate in the constrained model are extremely exxaggerated. This does not effect our estimation method directly, as we do not use or attempt to estimate the dispersal rate, but rather rely on patterns resulting from the partitioning of the dispersal rates between lineages. However this variance in effective global dispersal rate might lead to reduction of power in our method, as any differences due to how the dispersal is partition or distributed across various components of the system may be obscured. This variance in dispersal rates is due to the violation of the assumptions of the DEC model with respect to dispersal. As noted above, in the constrained dispersal regime we weight the dispersal of lineages associated with the open habitat trait twice the global dispersal rate, to maintain, on the average, the same dispersal flux as the unconstrained dispersal regime. As can be seen here, this has the desired effect. The noise that we are seeing is that the weighting factor of 2 is an expectation based on the stationary distribution of the habitat trait state. This expectation, however, is only an expectation, and the stochasticity of the trait transition process adds noise to the dispersal estimates, as, in any one (constrained dispersal regime) simulation a varying and unpredictable number of lineages are not dispersing. The variance of the trait transition process increases strongly as the trait transition rates get large, as seen in 3b. If we “correct” for the trait transition rate, as can be seen in figure 3d, the dispersal rates of both the constrained and unconstrained dispersal regimes are more concordant.

### Performance Over Parameter Space

Figure 4 shows the performance of the method over a broad range of parameter space. Figure 4a shows the performance faceted by the diversification parameters (birth rate, *b*, in rows, and extirpation rate (as a factor of birth rate), *e*, in columns), while figure 4b shows a unified summary. In both cases, the *x*-axis is the dispersal rate and the *y*-axis the trait transition rate, both in (log) units of the birth rate. The main metric each plot shows is the performance of our method at the locality in parameter space defined by that combination of process parameters: birth rate, extirpation rate, trait transition rate, and dispersal rate. This performance is measured in terms of the proportion of test data sets that were generated from that locality in parameter space that were correctly classified (in terms of the “unconstrained” vs. “constrained” dispersal regime) when using training data sets generated in the same locality in parameter space. As can be seen in the faceted plot, figure 4a, the general performance of the method is dominated with the *relative* birth rate rather than the absolute.

**Figure 4.**
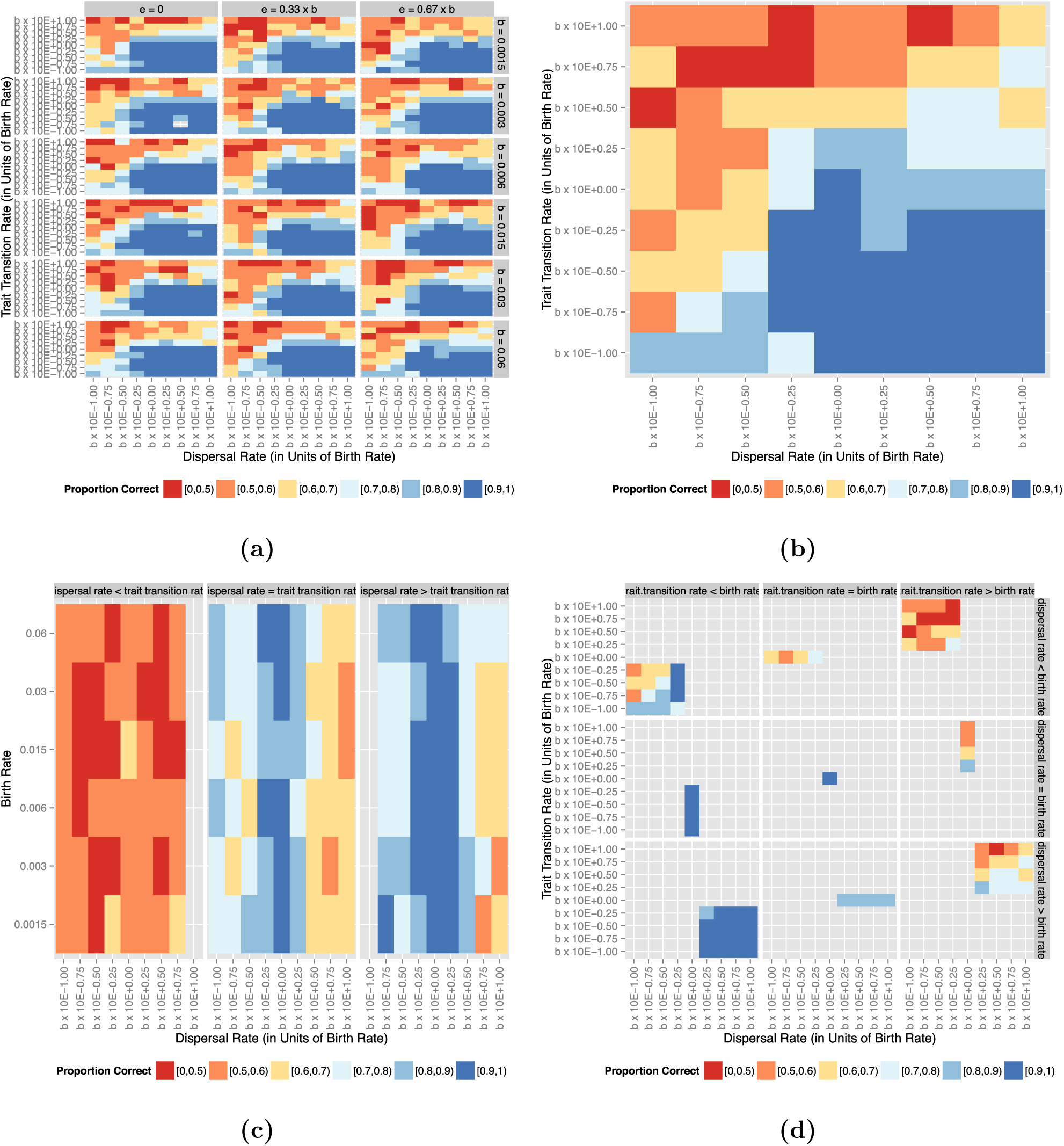
Performance of model-choice by simulation-trained discriminant analysis function classification over parameter space, as measured in proportion of correct classifications over 200 replicate analysis at each point in parameter space. Dispersal rates are on the *x*-axis and trait evolution transition rates are on the *y*-axis, expressed in units of birth rate (so that, for example, a value of “*b ×* 10E – 1.00” indicates a rate one order of magnitude less than the birth rate). Plots are (a) faceted to indicate different birth rates in rows and extirpation rates in columns, (b) marginalized over birth rates and extirpation rates, (c) partitioned into dispersal rates relative to trait transition rates, and (d) partitioned into dispersal rates relative to trait transition relative to birth rates.

Figure 4b shows the same parameter space, but reduces the visualization dimensionality by reducing the birth rate to a relative scaling factor and marginalizing over extirpation rates. What stands in this plot is that there is clear area where out method performs well and a clear area where it does not, and these areas can be defined in terms of the rates of trait evolution and dispersal relative to the birth rate as well as each other. What figure 4b shows is that, generally, the method performs best when the dispersal rate is *equal to or greater* than the trait transition rate. This becomes strikingly evident in figure 4c, which shows the differences in performance when the dispersal rate is less than, equal to, or greater than the trait transition rate. As before, the dispersal rate on the *x*-axis is in units of log birth rate. The birth rate is shown on the *y*-axis, and, as can be seen, the performance of the method is dominated by the relative values of the dispersal rate and the trait transition rate. Figure 4d shows the regionalization of parameter space in terms of performance at an even higher resolution, faceting the performance plots by the relative values of the transition rates to birth rate in colums, and the dispersal rates to birth rates in rows. What is seen here is that, in addition to the dispersal rates being higher than or equal to the trait transition rates, the optimum performance is gained when the birth rate, as well, is higher than or equal to the trait transition rate.

Figure 5 examines in detail the performance over parameter space as measured in terms of posterior probabilities found for the true vs. false models. This figure collapses the entire parameter space into a single axis, expressing it as the dispersal rate in units of trait evolution transition rate on the *x*-axis, with each point marginalizing over the various birth- and extirpation-rates. Thus, for example, the value indicated by “ × 10E–1.00” encompasses all points in parameter space where the dispersal rate is 0.01 the value of the trait evolution transition rate, over all different birth rates and extirpation rate. The distribution of posterior probabilities across all analyses for a particular point in parameter space is shown on the *y*-axis, with the color indicating the value range of the posterior probability and the relative span of the color indicating the relative proportion of analyses resulting in that posterior probability. The left column shows the posterior probability of support for the *false* model, while the right column shows the posterior probability of support for the *true* model; the top row shows analyses where the true model was “constrained” while the bottom row shows analyses where the true model was “unconstrained”. There are two important observations to be made here. First, that there is no difference in response between the “constrained” and “unconstrained” cases, i.e., the performance of the method is invariant with respect to the true model. The second, and perhaps for greater practical interest, is that the posterior of the false model very rarely reaches or exceeds 0.90, even in the suboptimal parts of parameter space where it is preferred over the true model. In constrast, the support for the true model generally exceeds 0.90, and often 0.99, in the optimal parts of parameter space.

**Figure 5.**
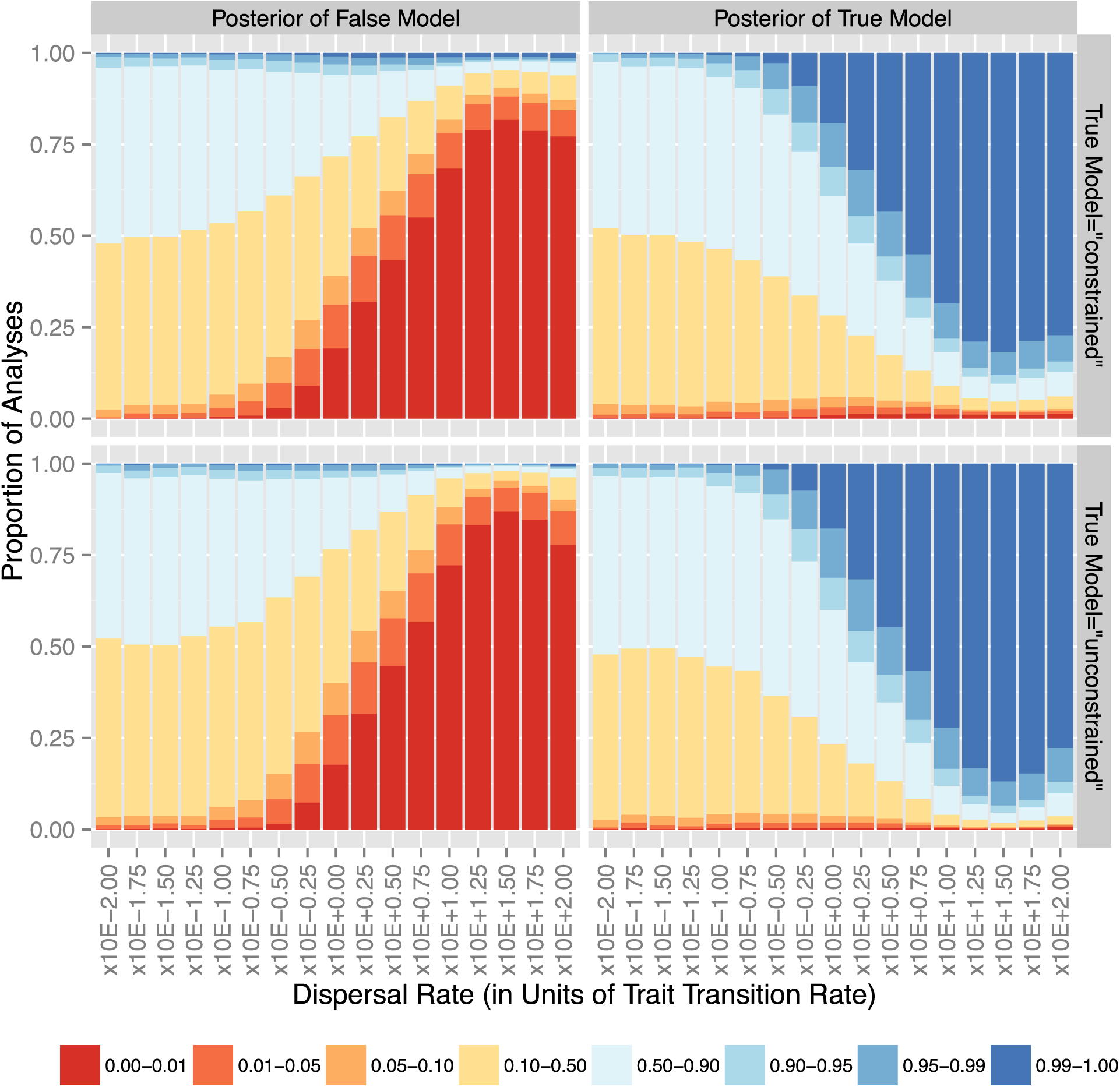
Performance over parameter space as measured by posterior probabilities given by the DAPC classification function of the true and false models. The cases where the true (generating) model corresponds to the “constrained” dispersal regime are shown in the first row, while the cases where the true model corresponds to the “unconstrained” dispersal regime are shown in the second row. The left column shows how support for the false model (i.e., the “unconstrained” dispersal regime if the true regime is “constrained”, and vice versa) varies over parameter space, while the right column shows the same for the *true* model. In each of the subplots, the *x*-axis shows dispersal rate in units of trait transition rate (so that, for example, a value of “ × 10E –1.00” indicates a dispersal rate one order of magnitude less than the trait transition rate). The *y*-axis shows the break-down of the multiple analyses carried out in that part of parameter space in terms of the different posteriors resulting for either the false or true model. The relative portion of the *y*-axis spanned by a sub-bar of a particular color indicates the proportion of the replicate analyses that yielded a posterior probability in favor of the false (on the left) or true (on the right) model corresponding to that color.

### Sensitivity to Calibration Errors

Figure 6 shows the effect of errors in the process calibration parameters on the posterior probability of the true model. As before, the *y*-axis shows the relative proportion of posterior probabilties across analyses for a particular set of parameters. The *x*-axis, on the other hand, shows the magnitude of error introduced in the birth, dispersal and trait evolution transition rates in the left, right, and center panels, respectively. In each panel, the set of analyses with *no* error in the calibration parameters is indicated by a vertical ine. We note that the method is sensitive to errors in birth rate parameter (show in the left-most panel) is “unconstrained”, and somewhat less so when the true model is “constrained”. We suggest, that, in the extreme cases, these results are an artifact of speciation occurring either too slow or too fast to generate a pattern of diversity (with distributions of traits and areas across lineages) for the summary statistics to capture. The method is also sensitive to error in the dispersal rate parameter, especially when the error is due to an underestimate in the dispersal rate and the true model is “unconstrained”. In both the “unconstrained” and “constrained” cases, the method is fairly robust to an overestimate of the dispersal rates, up to an order of magnitude in the former and two orders of magnitude in the latter. The method is least sensitive to error in the trait evolution transition rate parameter, with very little discernible reduction in performance over *two* orders of magnitude difference in rate, regardless of whether or not the true model is “unconstrained” or “constrained”.

**Figure 6.**
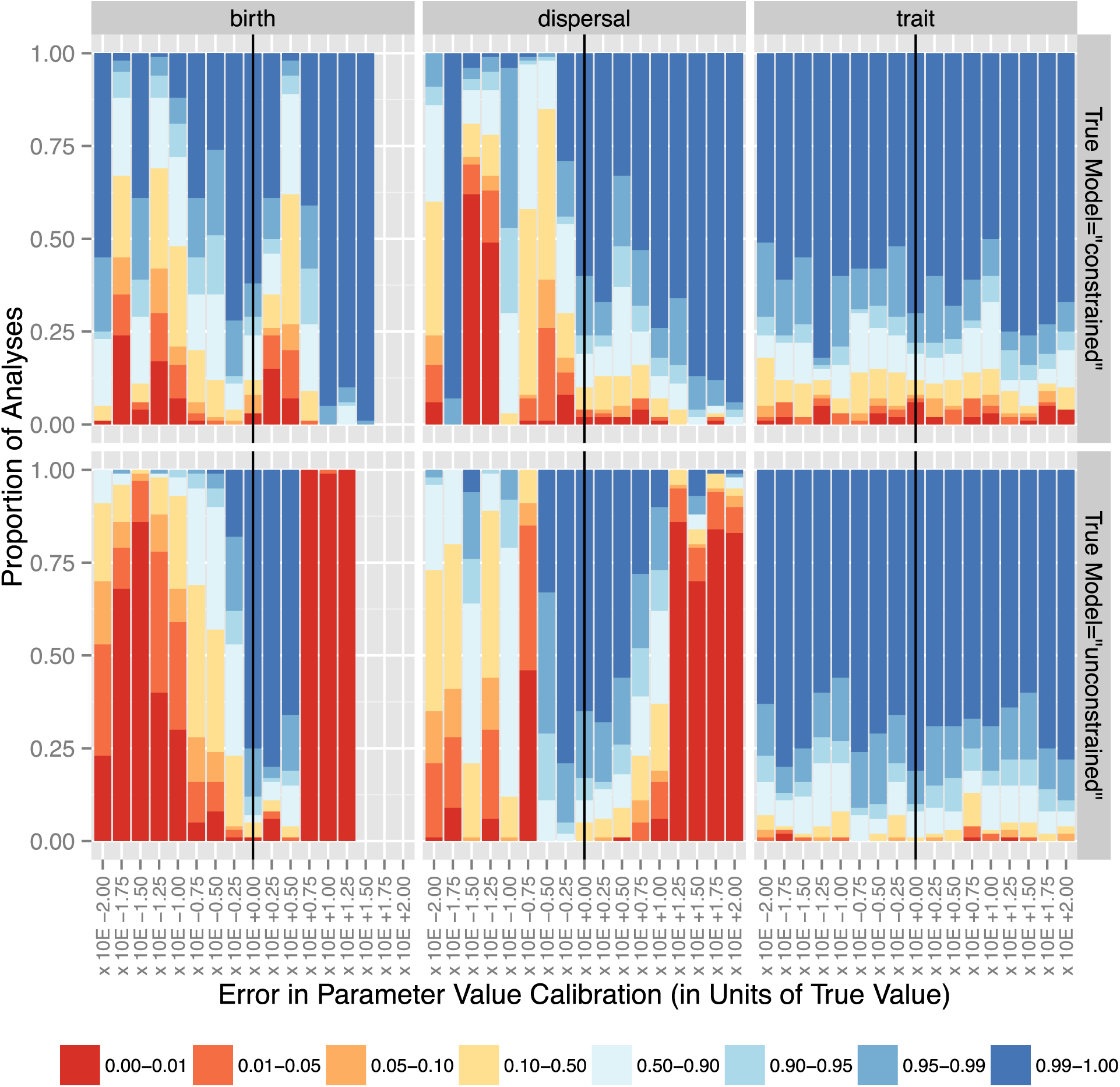
Changes in performance, as measured by posterior probability of support for the true model, as errors of different magnitudes are introduced into the training data set calibration values for the birth (left), dispersal (center), and trait evolution transition (right) rates. The magnitude of the error is indicated on the *x*-axis, with “*b ×* 10E+0.00” indicating *no* error (marked with the black vertical line). As in figure 5, above, the relative portion of the *y*-axis spanned by a sub-bar of a particular color indicates the proportion of the replicate analyses that yielded a posterior probability corresponding to that color.

### Dispersal in Island Birds

#### Habitat-Dependent Dispersal

The Wallacean bird system phylogeny of bird lineages categorized by their habitat trait had 365 species, with a maximum-likelihood estimate of the birth rate under a pure-birth model of *b* = 0.05482, and the maximum-likelihood estimates of the global dispersal rate and extirpation rate between all island modules under a DEC model of *d* = 0.01798 and *e* = 0.02579, respectively. The maximum-likelihood estimate of the open habitat vs. interior habitat trait transition rate under an equal-rates model was *q*_habitat_ = 0.047111. The dispersal rate is lower than the birth and trait transition rates, which, as the previous results indicate, might be in a sub-optimal region of parameter space.

The results for the discriminant analyses are shown in Table 1 and 2. Four separate analyses were carried out using training datasets generated under Archipelago models calibrated with the above process parameter values. The first three used zero, one, and two supplementary areas, while the last used a training data set that combined the previous three (i.e., with an equal mixture of zero, one, two and three supplementary areas.

**Table 1:** Assessment of discriminant analysis function efficacy for Wallacean bird fauna habitat-constrained dispersal classification by (re-)application of discriminant analysis function on training data sets.

| Sup-<br>ple-<br>men-<br>tary<br>Areas | Re-<br>tain-<br>ed<br>PC<br>Axes | Proportion<br>of Correct<br>Reclassifi-<br>cations | Mean<br>Poste-<br>rior of<br>True<br>Model | Proportion of Correct<br>Classifications When<br>True Model was<br>“Unconstrained” | Proportion of Correct<br>Classifications When<br>True Model was<br>“Constrained” | Proportion of<br>Incorrect<br>Classifications When<br>True Model was<br>“Unconstrained” | Proportion of<br>Incorrect<br>Classifications When<br>True Model was<br>“Constrained” |
| --- | --- | --- | --- | --- | --- | --- | --- |
| 0 | 31 | 0.83500 | 0.75657 | 0.49701 | 0.50299 | 0.51515 | 0.48485 |
| 1 | 40 | 0.91000 | 0.84867 | 0.50000 | 0.50000 | 0.50000 | 0.50000 |
| 2 | 34 | 0.88500 | 0.80032 | 0.50282 | 0.49718 | 0.47826 | 0.52174 |
| 0+1+2 | 40 | 0.73667 | 0.64296 | 0.50000 | 0.50000 | 0.50000 | 0.50000 |

Table 1 shows the profile of the discriminant analysis functions constructed on the four supplementary area configurations, and the results of applying each of these functions to classify the data in the training data set. As can be seen, the proportion of correct (re-)classifications ranges from 0.73, in the case of the union of all three training data sets, to 0.91, in the case of the a single supplemental area. Note that, as discussed in the Discussion section, it remains to be determined whether or not posterior probabilities can be compared across different analysis incorporating different models such as this case, and thus we caution that these result should *not* be taken to indicate that one supplemental area is in any way a preferred solution over others. More informative is that we see that there is very little bias in the method in most cases: both the correct classifications as well as the incorrect classifications are more or less equally distributed between the “constrained” and “constrained” models.

Table 2 shows the results of applying the discriminant analysis function on the empirical data. As can be seen, regardless of the number of supplemental areas used, all analyses strongly supported a “constrained” dispersal regime, i.e. that dispersal is indeed constrained to lineages associated with the open habitats, with a posterior probability that exceeds 0.99. Even though this analysis falls in a sub-optimum region of parameter space (*c.f.* figure 4, the very high posterior probability in support for the “constrained” model is encouraging, given the results shown in figure 5.

**Table 2:** Classification of Wallacean bird fauna with respect to whether dispersal is dependent on habitat type, using training data sets constructed with varying numbers of supplemental areas.

| Supplementary Areas | Retained PC Axes | Assigned Dispersal Regime | Posterior |
| --- | --- | --- | --- |
| 0 | 31 | constrained | 0.99418 |
| 1 | 40 | constrained | 0.99992 |
| 2 | 34 | constrained | 1.00000 |
| 0+1+2 | 40 | constrained | 0.99981 |

#### Trophic-Level Dependent Dispersal

The phylogeny of Wallacean bird lineages categorized by their trophic-level trait had 549 species, with a maximum-likelihood estimate of the birth rate under a pure-birth model of *b* = 0.062865, and the maximum-likelihood estimates of the global dispersal rate and extirpation rate between all island modules under a DEC model of *d* = 0.02258 and *e* = 0.003981, respectively. The maximum-likelihood estimate of the low to high trophic-level trait transition rate under an equal-rates model was *q*_trophic-level_ = 0.00537. These calibration parameters, with the dispersal rate much higher than the trait transition rate, place this set of analyses in what was determined to be an optimal part of parameter space.

Table 3 shows the profile of the discriminant analysis functions constructed on the four supplementary area configurations, and the results of applying each of these functions to classify the data in the training data set. Here, we saee that in the case of 0 supplemental areas, there is a stronger tendency to mis-classify the cases where the true mode was “constrained”, and this effect presumably influences the cases where the training data set used the union of the different supplemental area configurations. However, the analyses with one or two supplemental areas are unbiased with respect to the generating model, and the use of two supplemental areas has the peak proportion of correction (re-)classifications, with 0.91 replicates in the training data set correctly classified.

**Table 3:** Assessment of discriminant analysis function efficacy for Wallacean bird fauna trophic-level dependent dispersal classification by (re-)application of discriminant analysis function on training data sets.

| Sup-<br>ple-<br>men-<br>tary<br>Areas | Re-<br>tain-<br>ed<br>PC<br>Axes | Proportion<br>of Correct<br>Reclassifi-<br>cations | Mean<br>Poste-<br>rior of<br>True<br>Model | Proportion of Correct<br>Classifications When<br>True Model was<br>“Unconstrained” | Proportion of Correct<br>Classifications When<br>True Model was<br>“Constrained” | Proportion of<br>Incorrect<br>Classifications When<br>True Model was<br>“Unconstrained” | Proportion of<br>Incorrect<br>Classifications When<br>True Model was<br>“Constrained” |
| --- | --- | --- | --- | --- | --- | --- | --- |
| 0 | 26 | 0.84000 | 0.76132 | 0.52381 | 0.47619 | 0.37500 | 0.62500 |
| 1 | 26 | 0.84000 | 0.74755 | 0.50000 | 0.50000 | 0.50000 | 0.50000 |
| 2 | 23 | 0.91000 | 0.83198 | 0.50000 | 0.50000 | 0.50000 | 0.50000 |
| 0+1+2 | 27 | 0.81167 | 0.72967 | 0.50719 | 0.49281 | 0.46903 | 0.53097 |

Table 4 shows the results of applying the resulting discriminant analysis function on the empirical data. In constrast to the analyses of habitat-dependent dispersal, here all the analyses strongly support the *non*-constrained regimes: i.e., that trophic-level does *not* regulate or drive the dispersal process in Wallacean birds.

**Table 4:**
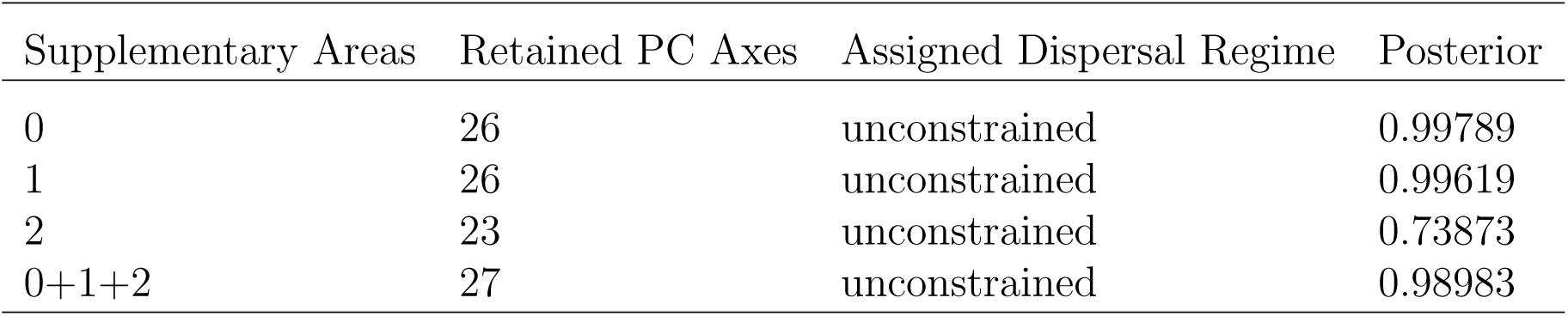
Classification of Wallacean bird fauna with respect to whether dispersal is dependent on trophic levels, using training data sets constructed with varying numbers of supplemental areas.

| Supplementary Areas | Retained PC Axes | Assigned Dispersal Regime | Posterior |
| --- | --- | --- | --- |
| 0 | 26 | unconstrained | 0.99789 |
| 1 | 26 | unconstrained | 0.99619 |
| 2 | 23 | unconstrained | 0.73873 |
| 0+1+2 | 27 | unconstrained | 0.98983 |

## DISCUSSION

### The “Archipelago” Model

The “Archipelago” model is an extremely flexible biogeographical phylogenesis model that not only simulataneously integrates the diversification, trait evolution, as well as the geographical evolution processes in a single unified superprocess, but also provides for these sub-processeses to inform each other dynamically as the phylogeny grows. This provides for the unique possibility of analyzing the influences of ecology, or any other biological trait, on *both* the diversification and geographical evolution process, in a way that is not possible using any other current biogeographical model.

Many biogeographical models, such as DEC or “`BayAreas`” etc., all assume exchangeable lineages, and cannot facilitate the exploration of the effects on lineage traits on either diversification or dispersal. GeoSSE (Goldberg et al. 2011) does allow for more sophisticated modeling of the diversifiction process by modeling the interaction between diversification rates and geographical range evolution However, as noted in the introduction, to study the effect of ecology on diversification, habitats are treated as geographic areas, which conflates ecology and geography, such that the interaction of ecology on geographical range cannot be studied with the GeoSSE approach.

In addition, unlike other existing methods, with our method, the ability to partition the geographical system into focal areas and supplemental areas, allowing the two classes of areas to interact biogeographically, yet only sampling data from the focal areas, solves a major conceptual limitation. Specifically, with community-based biogegoraphical analyses using previous approaches, the simulated data are typically generated under, e.g., a birth-death model, which results in a strict and complete monophyly of all sampled lineages: the entire diversity of the region is assumed to have evolved in-situ. However, if the study region exchanges fauna with external regions, then the simulated phylogenies fail to account for the fact that the empirical phylogeny is really paraphyletic with respect to the full diversity of the radiation unless there has only been a single colonization of the system. With some groups of limited scope, this latter case might indeed be true, but in most cases this is not.

### Simulation-Trained Discriminant Analysis of Principal Components Classification for Biogeographical Model Choice

We use a canonical discriminant analysis function trained by simulated data to classify a target data set, based on principal components of summary statistics calculated on both data sets. This method is extremely flexible, in that it can be used to carry out model selection with models for which likelihoods are too expensive to calculate in either an MCMC or maximum-likelihood optimization heuristc framework, or even cannot be calculated at all due to the difficulty in formulating a likelihood statement. In this respect, it is very similar to other simulation-based alternatives to full-likelihood inference, such as Approximate Bayesian Computation (ABC) approaches (e.g. Beaumont et al. 2002; Beaumont 2010; Marjoram et al. 2003). However, the posterior probabilities of the models in the approach we use are not directly comparable to the posterior probabilities of either full-likelihood Bayesian or ABC approaches for a number of reasons. For example, we develop the discriminant analysis functions based on simulations calibrated with process parameter values estimated from the original data, whereas both full-likelihood Bayesian and ABC approaches require the prior be specified without using the original data (though empirical Bayesian approaches use maximum-likehood estimates of nuisance parameters). Even without this issue, however, (e.g., if we were to use an independent data set to estimate these parameters), it is still not entirely clear if the posterior probabilities can be compared, due to a number of other factors in our approach that are difficult to relate to these other approaches; for e.g., the selection of the number of PC axes to retain in the construction of the DA function. As such, we prefer to view our approach as more akin to a machine-learning approach, i.e., model classification as opposed to model selection, which, indeed, strictly speaking it truly is. The posterior probabilities obtained by our approach might be taken to be the relative strength of support for different candidate models in the same pool (i.e., within the same single discriminant analysis), but at this point more work needs to be done to establish how to compare these values across different analysis or the posterior probabilities produced by full-likelihood or ABC approaches.

We cannot use full-likelihood approach to estimate under all but the simplest case of our model (e.g., where rates of speciation, dispersal, and so on are not dynamically-informed by the states of lineages, but are instead fixed to be true homogenous Poisson processes), but why do we not use ABC? The primary reason is that summary statistic development in ABC is very complex, requiring detailed exploration of parameter space to establish the efficacy of the summary statistics with respect to the particular configuration of model space and data (Robert et al. 2011). Approximate Bayesian Computation has been shown to be a fairly robust procedure for parameter estimation, but, it has been shown to have many issues with model selection, and it takes an immense amount of effort to develop summary statistics that are effective for a particular model pool, and even more effort to *demonstrate* this effectiveness and to understand and characterize the biases that might arise with any particular suite of summary statistics (e.g. Oaks et al. 2013, 2014). While we do conduct such a detailed exploration here, in principle with the DAPC approach, this is not neccessary in practice for any particular study: re-application of the discriminant analysis classification on the training data set allows immediate evaluation of the efficacy of the summary statistics used in terms of the proportion of elements in the training data that were classified correctly (supplemented by, if preferred, the mean posterior probability of the true model), as well as the biases that might be apparent (e.g., if one model is preferred more than any other when mis-classified). Furthermore, with ABC approaches, selection of summary statistics is always a balance between not having too many summary statistics such that it becomes difficult to simulate data that falls within the acceptence threshold, or having too few such that resolving power is lost. With the DAPC approach used here in conjunction with the heuristic we developed to select the number of principal component axis to retain, this issue is largely alleviated. Another reason to use the DAPC approach over the ABC is due to computational efficiency. An ABC analysis requires a large number of simulations to adequately sample parameter space, unless these are fixed or reduced by using empirically-informed priors, though this latter needs to be used with caution (Oaks et al. 2014). As shown here, effective results are available with our approach with as few as 200 simulation replicates.

There are four major limitations with our approach to model choice via simulation-trained discriminant analysis classification, at least as presented here.

The first is that, while immensely more tractable than ABC, our approach still requires formulation and testing of summary statistics. The summary statistics that we use here are explicitly design to capture differences in the patterns of how areas and traits sort themselves on a phylogeny and are partitioned with respect to each other. This is suitable for the types of studies presented here, i.e., the relationship between dispersal and traits, but may not be for the broader classes of studies that might be carried out with the “Archipelago” model. As noted above, however, this task is greatly facilitated by the ease and effectiveness of summary statistic assessment through reapplication of the result discriminant analysis function on the training data.

The second and third issues are related: the need for external estimation of parameters, and treating of uncertainty in these estimates. As presented here, the speciation, extinction, dispersal, and trait transition rate parameters of the model need to be calibrated using estimates made on the empirical data. This is less than ideal both for theoretical as well as practical reasons. The theoretical objection is related to the same objection expressed against empirical Bayesian approaches, i.e., in some sense the data is used “twice”. We assert that this is not a salient concern in our case, as we are not trying to recover, even approximately, posterior probabilities comparable to Bayesian posterior probabilities, but are rather focussing on model choice. The practical concern is that the dispersal parameter, in particular, is computationally too intensive to estimate once the number of areas gets very large using either “`BioGeoBears`” (Matzke 2014) or “`lagrange`” (Ree and Smith 2008), especially since, in the types of studies we present here, we allow any possible combination of areas for the ranges of internal nodes. This means that while, in principle, our approach can easily handle relative large numbers of areas with still very reasonable analysis times (including simulation and summary statistic calculation), the need to estimate the calibration parameters using more traditional approaches limits the number of areas to those that can be handled by either of the aforementioned packages.

Furthermore, even if the parameters can be obtained readily from the empirical or target data to calibrate the training data simulations, we do not treat the uncertainty in these estimates in the approach that we present hee. To some extant, it can be argued that the “fuzziness” of our approach allows for a large degree of slop in the calibration parameters, and, indeed, we demonstrated this in our results, where, depending on the parameters, errors of moderate to even large degrees of magnitude can be tolerated. One modification to our approach that allows both for treatment of uncertainty as well as avoiding the need to rely on other methods to estimate the calibration parameters is possible, albeit at an obvious and substantial increase in computational cost. Basically, the training data sets need to be constructed over a broad range of parameter space, with the parameter values of each training data element being sampled randomly from a prior distribution, thus both obviating the need for external estimates of calibration parameters as well as integrating out uncertainty in these parameter values. This brings our approach closer to ABC operationally, though theoretically both a very distinct, and operationaly the two remain different in how the model support is assessed. For example, in our empirical application presented here, the fourth class of training data set, i.e. the one that combined zero, one, and two supplementary areas, can be seen as implementing this concept to integrate out uncertainty in the number of supplemental or external areas that might be interacting with the focal areas under a prior that is uniform over the range 𝒰 {0, 2}.

Like all model-selection approaches, full-likelhood Bayesian or otherwise, the analysis is unable to choose a model that it is not offered, and thus, in a sense, any analysis result can only be as “true” as the models that are constructed by the researcher. And we assert that this is not a problem, or a limitation, but rather the basis of model-based science. While model-*selection* or choice should be carried out through rigorous and objective statistical procedures, model *formulation* is, we suggest, the primary task of the investigator, *not* the statistical apparatus. Formulating a model is the way for investigators to express their ideas, expertise, knowledge to a statistical framework. As the first step in an analysis, it is essentially equivalent to “asking” the question, and while statistics and data have the capacity to answer questions, asking the question remains and should always remain the role of the investigator.

### Conclusions

Biogeographic dynamics involve the interplay of speciation, extinction, and dispersal in a historical context of climate and geologic change, but in lieu of direct observations and experimental paradigms, inferring such dynamics is a major quantitative challenge. Here we introduce a model coupled with an inference method that allows for the study of the relationships and interaction between these processes, as given by the patterns evident in the phylogeny that they generate over evolutionary history. This is the first time these biogeographical and evolutionary processes, so fundamental and central to a large field of biogeographical inquiry and theory, have been explicitly integrated in phylogeny-based model choice framework. While we only explore the relationship between habitats and dispersal in detail here, it should be obvious that the model and method are flexible enough to consider various other types of analyses, where ecology and geography inform not only each other, but also the diversification process.

The flexibility and conceputal breadth of our model is coupled with an inference method that is not only computationally efficient (requiring only 200 simulation replicates to generate effective training datasets across a broad range of parameter space), but also effective over a broad range of parameter space that is important for many systems. Using our method, we demonstrate a strong support for dispersal in Wallacean birds being regulated by their habitat traits, with dispersal being constrained to lineages associated with open or marginal habitats, in accord with previous statistical work on the group (Carstensen and Olesen 2009; Carstensen et al. 2013). Such constraints are the building blocks of larger integrative theories such as the taxon cycle, and as such we feel our method is a step toward the broader goal of developing a quantitative basis for testing complex biogeographic theory.

## ACKNOWLEDGEMENTS

This work was supported by subsidy funding from OIST and an NSF grant (DEB-1145989) to EPE and LLK. We also wish to thank Mark T. Holder at the University of Kansas for his generosity with the HPC computational resources of his lab, without which this work would not have been possible. All plots were done using the “`R`” package, “`ggplot2`” (Wickham 2009).

